# Axonal CB1 receptors mediate inhibitory bouton formation via cAMP increase

**DOI:** 10.1101/2021.04.12.439503

**Authors:** Jian Liang, Dennis LH Kruijssen, Aniek CJ Verschuuren, Bas JB Voesenek, Feline Benavides, Maria Sáez Gonzalez, Marvin Ruiter, Corette J Wierenga

## Abstract

Experience-dependent formation and removal of synapses are essential throughout life. For instance, GABAergic synapses are removed to facilitate learning, and strong excitatory activity is accompanied by formation of inhibitory synapses. We recently discovered that active dendrites trigger the growth of inhibitory synapses via CB1 receptor-mediated endocannabinoid signaling, but the underlying mechanism remained unclear. Using two-photon microscopy to monitor the formation of individual inhibitory boutons, we found that CB1 receptor activation mediated formation of inhibitory boutons and promoted their subsequent stabilization. Inhibitory bouton formation did not require neuronal activity and was independent of G_i/o_ protein signaling, but was directly induced by elevating cAMP levels using forskolin and by activating G_s_ proteins using DREADDs. Our findings reveal that axonal CB1 receptors signal via unconventional downstream pathways and that inhibitory bouton formation is triggered by an increase in axonal cAMP levels. Our results demonstrate a novel role for axonal CB1 receptors in axon-specific, and context-dependent, inhibitory synapse formation.

## Introduction

Synaptic plasticity, the strengthening and weakening of existing synapses, is often considered the physiological basis for learning and adaptation. In addition, the experience-dependent formation and removal of synapses is equally important (Bailey and Kandel, 1993; Caroni et al., 2012). Changes in the number of synaptic connections have been shown to be critical during learning *in vivo* (Bailey and Chen, 1989; Caroni et al., 2012; Hofer et al., 2009; Kozorovitskiy et al., 2012; Ruediger et al., 2011) and strongly determine postsynaptic function (Scholl et al., 2020). Plasticity of GABAergic synapses is particularly important for shaping and controlling brain activity throughout life (Chiu et al., 2019; Flores and Méndez, 2014; Herstel and Wierenga, 2021; Maffei et al., 2017) and GABAergic dysfunction is associated with multiple brain disorders, including schizophrenia and autism (Lewis et al., 2005; Mullins et al., 2016; Tang et al., 2021). For example, the number of inhibitory synapses is rapidly adjusted during learning (Bourne and Harris, 2011; Chen et al., 2015; Donato et al., 2015, 2013) or when sensory input is lost (Keck et al., 2011) to facilitate plasticity at nearby excitatory synapses. Vice versa, potentiation of excitatory synapses can trigger the formation of inhibitory synapses to maintain a local balance (Bourne and Harris, 2011; Hu et al., 2019; Knott et al., 2002). The formation, stabilization and removal of synapses likely requires local context-dependent signaling mechanisms (Hu et al., 2019; Kirchner and Gjorgjieva, 2019; Kleindienst et al., 2011; Niculescu et al., 2018; Nishiyama and Yasuda, 2015; Oh et al., 2016), but our current understanding of these processes, especially at inhibitory synapses, is far from complete.

We recently discovered that strong, clustered activation of excitatory synapses along dendrites of hippocampal CA1 pyramidal neurons can trigger the formation of a new inhibitory bouton onto the activated dendrite (Hu et al., 2019). We proposed that this dendritic mechanism serves to maintain local balance between excitatory and inhibitory inputs during ongoing synaptic plasticity. Inhibitory bouton formation required dendritic endocannabinoid synthesis and activation of CB1 receptors (Hu et al., 2019). Dendritic endocannabinoids are well-known to serve as retrograde signals to regulate synaptic plasticity (Alger, 2002; Castillo et al., 2012; Chevaleyre and Castillo, 2003; Kano et al., 2009; Katona and Freund, 2012), but it is unclear how CB1 receptors can trigger new inhibitory bouton formation.

CB1 receptors are G-protein coupled receptors and are widely abundant in the brain. They are expressed in both excitatory and inhibitory neurons, as well as in glia cells (Bonilla-Del Río et al., 2021; Hebert-Chatelain et al., 2016; Maroso et al., 2016; Navarrete et al., 2014). Perhaps the most prominent CB1 expression is in a subset of inhibitory axons in the dendritic layer of the hippocampal CA1 area (Bonilla-Del Río et al., 2021; Dudok et al., 2015). Axonal CB1 signaling plays an important role during axon guidance (Argaw et al., 2011; Berghuis et al., 2007; Njoo et al., 2015; Roland et al., 2014), but axonal CB1 receptor expression remains high during adulthood. The best described actions of CB1 receptors in adulthood is to suppress neurotransmitter release (Alger, 2002; Castillo et al., 2012; Kano et al., 2009). However, CB1 receptors are not enriched in boutons, but freely diffuse within the entire axonal membrane (Dudok et al., 2015). It is possible that axonal CB1 receptors may function as replacement pool for internalized synaptic receptors at boutons as recently suggested for opioid receptors (Jullié et al., 2020), although synaptic enrichment would still be expected. In addition, GABA release at dendritic inhibitory synapses is not strongly modulated by CB1 receptors (Lee et al., 2015, 2010), and coupling between CB1 receptors and the active zone is weak (Dudok et al., 2015). This suggests that CB1 receptors in inhibitory axons serve an additional purpose. Interestingly, it was recently described that CB1 receptors can also mediate synaptic potentiation (Cui et al., 2016; Monday and Castillo, 2017; Wang et al., 2017). Although CB1 receptors typically signal via G_i/o_-proteins, many additional downstream pathways, both dependent and independent of G-proteins, have been described (Berghuis et al., 2007; Cui et al., 2016; Flores-Otero et al., 2014; Glass and Felder, 1997; Roland et al., 2014; Zhou et al., 2019).

Here, we demonstrate that activation of axonal CB1 receptors can trigger the initial formation of inhibitory synapses. Using two-photon time lapse imaging we observed the formation of inhibitory boutons upon brief application of the CB1 receptor agonist WIN. We demonstrate that this requires the presence of CB1 receptors on inhibitory axons. Furthermore, we found that CB1-mediated inhibitory bouton formation is independent of G_i/o_ protein signaling and neuronal activity. We find that new inhibitory boutons are formed in response to elevated cAMP levels or activation of G_s_ protein signaling in inhibitory axons. Our data indicate that activation of axonal CB1 receptors triggers inhibitory synapse formation via an atypical signaling pathway via G_s_-proteins. Furthermore, our data identify an increase in axonal cAMP as a crucial second messenger for mediating inhibitory bouton formation.

## Results

### Repeated CB1 receptor activation increases functional presynaptic terminals

We previously demonstrated that new inhibitory boutons can form in response to brief CB1 receptor activation (Hu et al., 2019). Newly formed boutons often did not persist (Hu et al., 2019), suggesting that additional or repeated signaling is required to eventually form functional inhibitory synapses (Frias et al., 2019; Wierenga, 2017). It was recently reported that strong, but brief, CB1 receptor activation can induce synaptic potentiation, while longer CB1 activation induces synaptic depression (Cui et al., 2016, 2015). This suggests that CB1 activation pattern is an important factor in determining its downstream signaling. We therefore sought to employ repeated, short activation of CB1 receptors in order to induce the formation of inhibitory synapses. We activated CB1 receptors in hippocampal slice cultures by repeated short exposure to the CB1 receptor ligand 2-AG (100 μM; 3 times 20 minutes with 2 hours interval) (Fig. 1A). We recorded miniature inhibitory postsynaptic currents (mIPSCs) in CA1 pyramidal neurons to assess functional inhibitory synapses 24 hours after the start of the first 2-AG exposure (Fig. 1B). Repeated CB1 receptor activation resulted in an increase of the mean mIPSC frequency by 38% (control: 3.9 ± 0.3 Hz; 2-AG: 5.5 ± 0.4 Hz, p=0.013; Fig 1C), while mIPSC amplitudes were not affected (Fig. 1D). Continuous exposure to 2-AG for 24 hours did not alter frequency or amplitude of spontaneous IPSCs (Fig. 1E,F), consistent with the notion that activation pattern determines CB1 downstream signaling. Interestingly, mIPSCs after repeated 2-AG exposure appeared to have longer rise times (Fig. 1G), while decay times were not different (Fig. 1H). We separated mIPSCs with slow and fast rise times based on a double Gaussian fit of the distribution of rise times (Fig. 1I). When we then analyzed the interevent intervals of fast and slow mIPSCs separately, we observed that the interevent intervals of slow mIPSCs were decreased after repeated CB1 activation, while the interevent intervals of fast mIPSCs were not affected (Fig. 1J,K). This analysis revealed that the observed increase in mIPSC frequency was due to a specific increase in the frequency of slow mIPSCs with long rise times (Fig. 1L). The rise time of mIPSCs depends on synaptic maturation (Gonzalez-Burgos et al., 2015; Lazarus and Josh Huang, 2011; Pardo et al., 2018), but is also strongly influenced by subcellular location, as dendritic filtering attenuates mIPSCs originating from dendritic inhibitory synapses (Bekkers and Clements, 1999; Rall, 1967; Wierenga and Wadman, 1999). This suggests that the increased mIPSC frequency after CB1 receptor activation may reflect an increase of inhibitory currents from dendritic locations, or from immature synapses.

**Figure 1.**
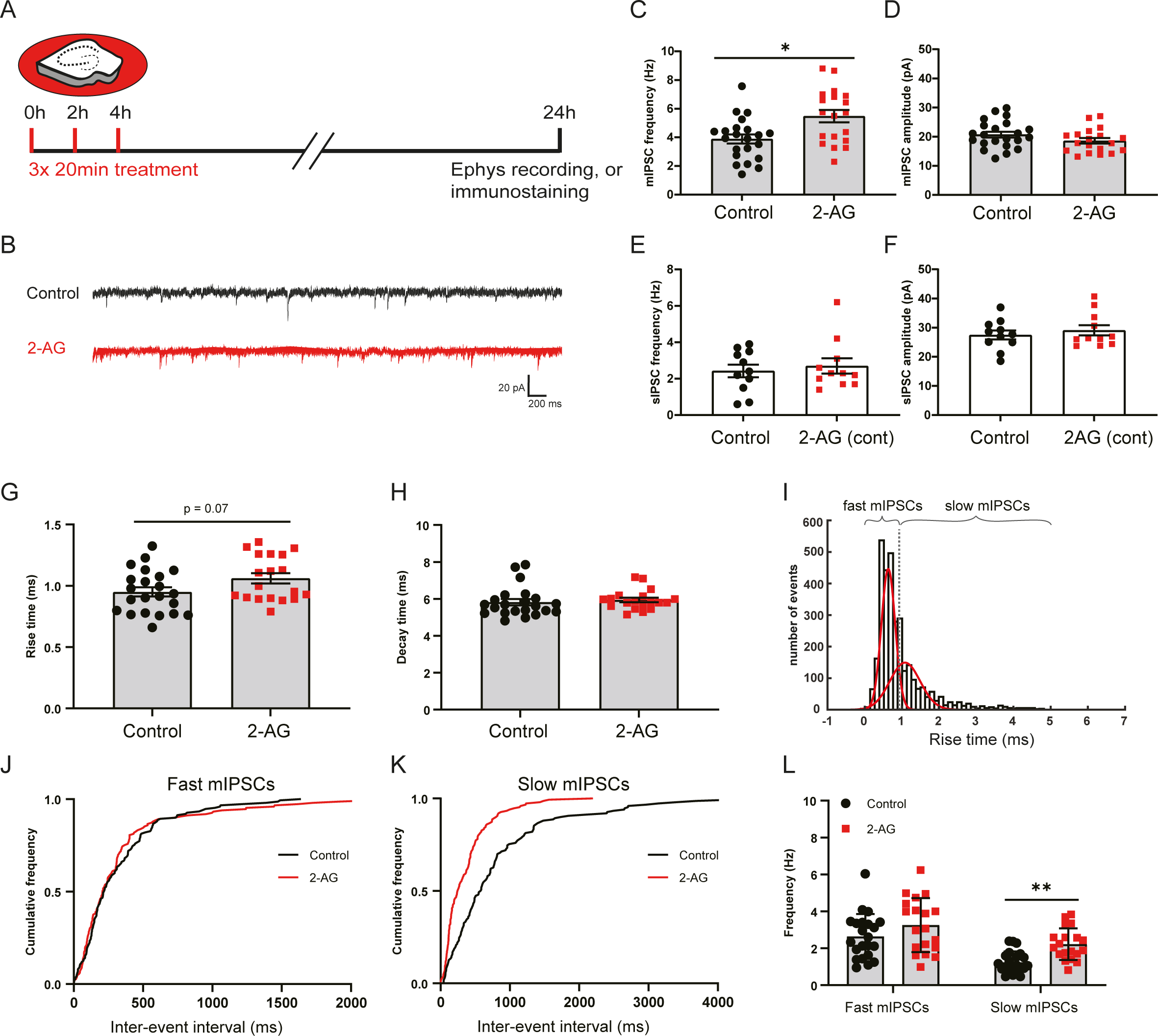
Repeated CB1 receptor activation results in increased mIPSC frequency. **(A)** Organotypic hippocampal cultures were treated 3 times with culturing medium containing 100 µM 2-AG or DMSO (control) for 20 minutes with 2 hour intervals. After 24 hours, slices were used for electrophysiology and immunostaining experiments. **(B)** Example traces of miniature inhibitory postsynaptic currents (mIPSCs) recordings from control (black) and 2-AG treated slice (red). **(C, D)** Mean frequency (C) and amplitude (D) of mIPSCs in control and 2-AG treated slices (MW, *p* = 0.013 in C and *p* = 0.16 in D). Data from 22 cells in 6 control slices and 19 cells in 6 2-AG treated slices. **(E, F)** Mean frequency (E) and amplitude (F) of sIPSCs in control and 2-AG treated slices, when 2-AG was continuously present for 24 hr (*p* = 0.99 in E and *p* = 0.95 in F; MW). Data from 11 cells in 5 control slices and 11 cells in 6 2-AG treated slices. **(G)** Mean rise time of mIPSCs in control and 2-AG treated slices (MW, *p* = 0.073). **(H)** Mean of mIPSC decay time in control and 2-AG treated slices (MW, *p* = 0.19). **(I)** The distribution of rise times of mIPSCs was fitted with a double Gaussian to separate fast and slow mIPSCs. **(J,K)** Cumulative distribution of interevent intervals of mIPSCs with fast (J) and slow (K) rise times (KS, *p* = 0.65 in J, and *p* < 0.0001 in K). (L) Mean frequency of mIPSCs with fast and slow rise times (2w ANOVA Sidak, fast: *p* = 0.14; slow: *p* = 0.0095). Data in G-L and C,D are from the same data set.

To determine if the observed increase in mIPSCs was associated with an increase in the number of inhibitory synapses, we analyzed presynaptic VGAT and postsynaptic gephyrin puncta in the dendritic region of the CA1 area in parallel immunohistochemistry experiments (Fig. 2A). We observed that the density of VGAT puncta was slightly increased after repeated 2-AG application (Fig. 2B), while the VGAT puncta size was decreased (Fig. 2C). Gephyrin puncta density and size were not affected by repeated 2-AG exposure (Fig. 2D,E), and the density of inhibitory synapses, defined as VGAT-gephyrin associations, was also not different from control slices (Fig. 2F,G). We therefore made a distinction between VGAT puncta that were associated with gephyrin and VGAT puncta without gephyrin (Fig. 2A, last panel). We observed that the increase in VGAT density was due to a specific increase in VGAT puncta that were not associated with gephyrin (Fig. 2H). In contrast, the reduction in VGAT puncta size was mostly due to a reduction in size of VGAT puncta with gephyrin association (Fig. 2I). This suggests that repeated short activation of CB1 receptors has two separable effect on inhibitory synapses: on the one hand it leads to shrinkage of VGAT clusters at inhibitory synapses, possibly reflecting synaptic depression (Monday et al., 2020), while at the same time new VGAT clusters are formed which are not associated with the postsynaptic scaffold gephyrin. Live imaging experiments have shown that VGAT is rapidly recruited when new boutons are formed in inhibitory axons, and that gephyrin normally follows within a few hours (Dobie and Craig, 2011; Frias et al., 2019; Wierenga et al., 2008). Our data suggest that repeated CB1 receptor activation induces the formation of presynaptic VGAT clusters, likely reflecting immature inhibitory synapses.

**Figure 2.**
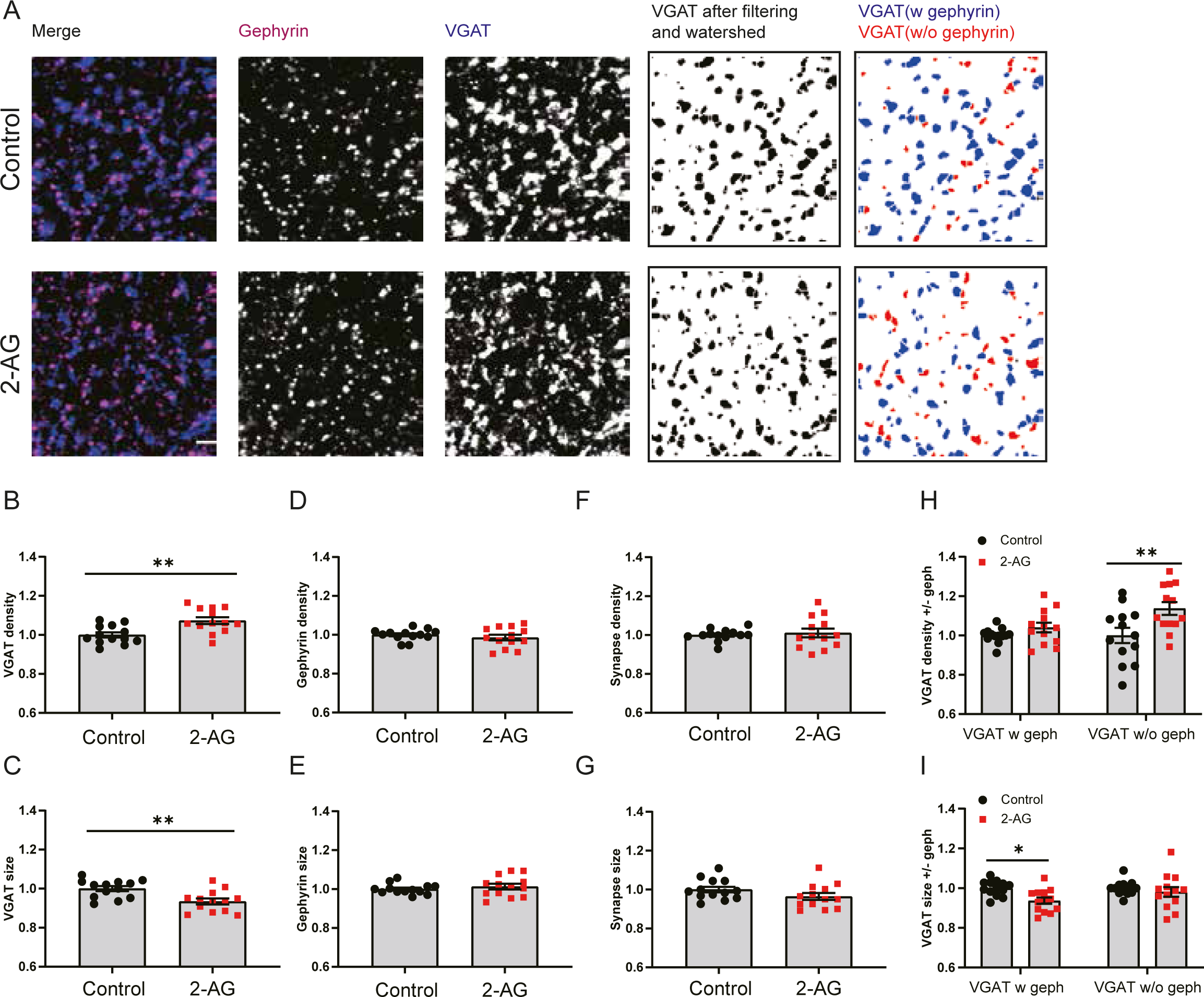
Repeated CB1 receptor activation induces the formation of partial inhibitory synapses. **(A)** Representative immunostaining images showing the presynaptic VGAT (blue) and postsynaptic gephyrin (purple) in control (upper) and 2-AG (lower) slices. Individual VGAT puncta were identified using watershed segmentation and these were color coded to distinguish VGAT puncta associated with gephyrin (blue) and VGAT puncta without gephyrin (red). **(B, C)** Normalized density (B) and size (C) of VGAT puncta in control and 2-AG slices (MW, *p* = 0. 0061 in B; *p* = 0.004 in C). **(D, E)** Normalized density (D) and size (E) of gephyrin puncta in control and 2-AG slices (MW, *p* = 0.54 in D; *p* = 0. 64 in E). **(F, G)** Normalized density (F) and size (G) of VGAT/gephyrin colocalizations in control and 2-AG slices (MW, *p* = 0.76 in F; *p* = 0.099 in G). **(H, I)** Normalized density (F) and size (G) of VGAT puncta with and without gephyrin (2w ANOVA Sidak, *p* = 0.55 and *p* = 0.003 in H; *p* = 0.017 and *p* = 0.65 in I). Data from 13 image stacks in 7 slices per group.

### Acute activation of CB1 receptors affects non-persistent boutons density only slightly

To get further insight in the role of CB1 receptors in the formation of inhibitory synapses, we performed two-photon live imaging in organotypic hippocampal slices to monitor GFP-labeled inhibitory bouton dynamics in response to short activation of CB1 receptors. Here we used short applications (5 minutes) of CB1 receptor agonists to mimic retrograde endocannabinoid signaling (Hu et al., 2019), but we wanted to avoid inducing synaptic weakening (Monday et al., 2020). We used the endogenous CB1 receptor ligand 2-AG as well as the chemically synthesized agonist WIN552121-2 (WIN), which is widely used because of its high affinity and stability (Chevaleyre et al., 2007; Roland et al., 2014; Wang et al., 2017). We verified that brief WIN application only transiently and mildly suppressed inhibitory currents (data not shown). As previously reported (Frias et al., 2019), the majority of inhibitory boutons were present at all timepoints during the 140 minutes imaging period (persistent boutons), but a substantial fraction of inhibitory boutons appeared, disappeared, or reappeared, during the imaging period (Fig 3A) (Frias et al., 2019; Schuemann et al., 2013). We will refer to the latter as non-persistent (NP) boutons. Bath application of 100 µM 2-AG (5 minutes) did not affect overall bouton density (control: 30.8 ± 1.7 boutons per 100 µm; 2-AG: 29.8 ± 1.7 boutons per 100 μm, p=0.81). The density of NP boutons appeared slightly increased after 2-AG compared to DMSO control (Fig. 3B,C), but this was mainly due to a large effect in a single axon. We calculated for each axon the average fraction of NP boutons that are present over time (NP presence). In control slices there was a small decrease in NP presence over time, possibly reflecting a decrease in network activity level when the slices are transferred from the incubator to the microscope. After 2-AG application NP presence appeared slightly more stable (Fig. 3D), but this difference did not reach statistical significance. We assessed if this difference could be traced back to a more specific effect in a particular subgroup of NP boutons (see methods and (Frias et al., 2019)), but we could not detect any differences in the densities of NP bouton subgroups in slices treated with control DMSO or 2-AG (Fig. 3E). There was also no difference in bouton duration (data not shown).

**Figure 3.**
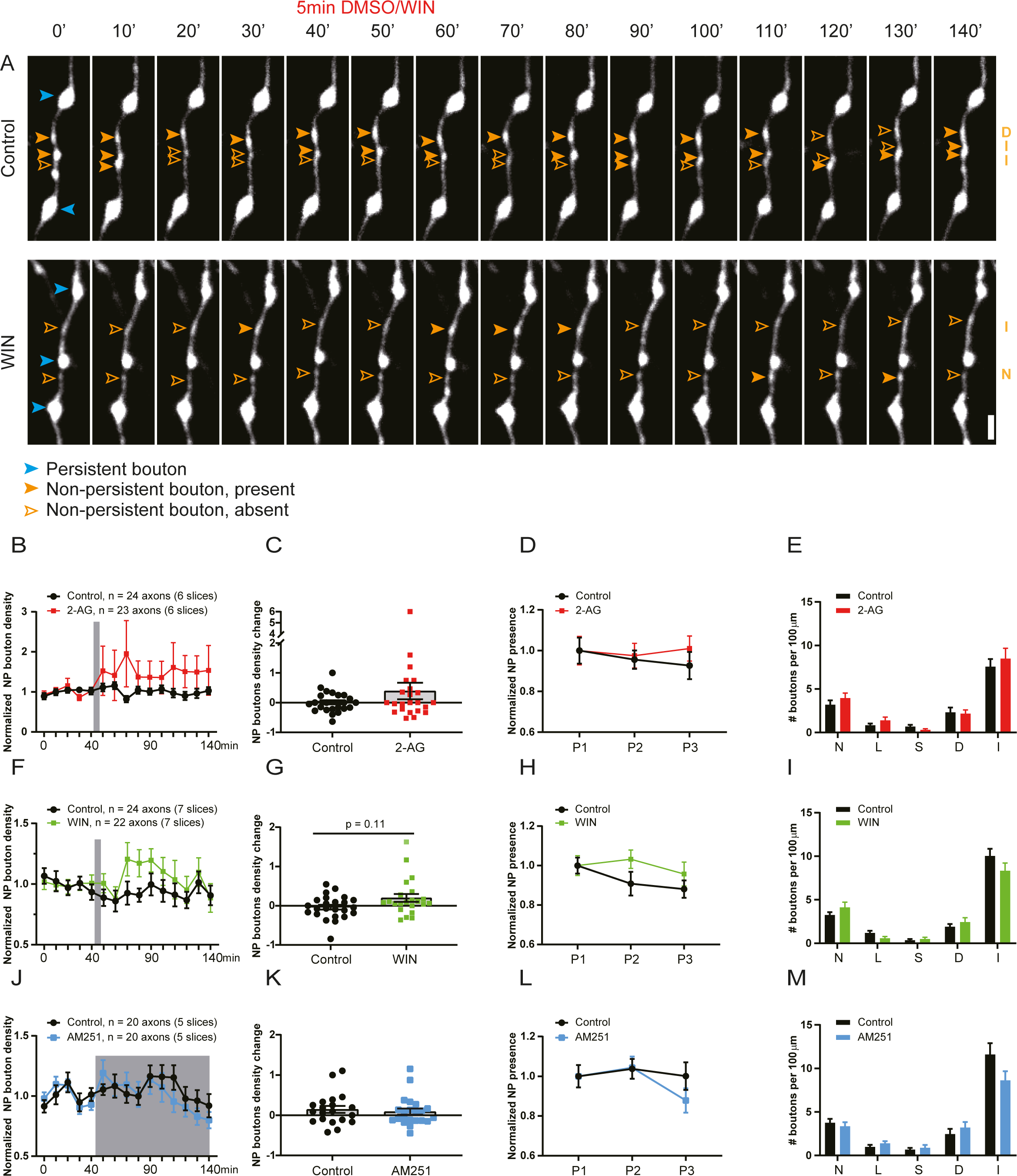
Brief activation of CB1 receptors slightly increases NP bouton density. **(A)** Representative two-photon time lapse images of GAD65-GFP labelled inhibitory axons in the dendritic region of the hippocampal CA1 area (maximal projections of 17 z-sections). After a baseline of five time points (40 minutes), CB1 receptor agonist or DMSO was washed in for 5 minutes. Imaging was continued for another ten time points (total imaging period is 140 minutes). Persistent boutons (blue) and non-persistent (NP) boutons (orange) are indicated by arrow heads. Empty arrow heads reflect a NP bouton which was absent at the time point. Scale bar is 2 µm. **(B)** CB1 receptors were activated by bath application of 100 μM 2-AG for 5 minutes. Normalized NP bouton density over time in control (black) slices and after 2-AG (red) application (2w ANOVA, *p* = 0.33). **(C)** Maximum change in NP bouton density in control slices and after 2-AG application (MW, *p* = 0.54). **(D)** Normalized NP presence over time in control and 2-AG treated slices. P1= time points 1 to 5, P2= time points 6 to 10, and P3= time points 11 to 15) in control and 2-AG treated slices (2w ANOVA, *p* = 0.61). **(E)** Mean density of NP bouton subgroups in control slices and after 2-AG application. N – new boutons (MW, *p* = 0.35); L – lost boutons (MW, *p* = 0.44); S – stabilizing boutons (MW, *p* = 0.21); D – destabilizing boutons (MW, *p* = 0.91); I – intermittent boutons (MW, *p* = 0.87). **(F)** CB1 receptors were activated by bath application of 20 μM WIN for 5 minutes. Normalized NP bouton density over time in control (black) slices and after 2-AG (green) application (2w ANOVA, *p* = 0.20). **(G)** Maximum change in NP bouton density in control slices and after WIN application (MW, *p* = 0.11). **(H)** Normalized NP presence over time in control slices and after WIN application (2w ANOVA, *p* = 0.20). **(I)** Mean density of NP bouton subgroups in control slices and after WIN application (MW, *p* = 0.40 (N); *p* = 0.06 (L); *p* = 0.79 (S); *p* = 0.70 (D); *p* = 0.10(I)). **(J)** Slices were treated with the CB1 receptor antagonist AM251 (5 μM) after time point 5. Normalized NP bouton density over time in control (black) slices and during AM251 (blue) application (2w ANOVA, *p* = 0.66). **(K)** Maximum change in NP bouton density in control slices and during AM251 application (MW, *p* = 0.6). **(L)** Normalized NP presence over time in control slices and during AM251 application (2w ANOVA, *p* = 0.56). **(M)** Mean density of NP bouton subgroups in control slices and during AM251 application (MW, *p* = 0.46 (N); *p* = 0.23 (L); *p* = 0.94 (S); *p* = 0.29 (D); *p* = 0.10(I)). Data in A from 24 axons in 6 control slices and 23 axons in 6 2-AG slices. Data in B from 24 axons in 7 control slices and 22 axons in 7 WIN slices. Data in C from 20 axons in 5 control slices and 20 axons in 5 AM251 slices.

The endocannabinoid 2-AG is rather unstable in solution and gets rapidly degraded in biological tissue (Dócs et al., 2017; Savinainen et al., 2012). To exclude the possibility that 2-AG gets degraded before it can activate CB1 receptors, we repeated these experiments using 20 µM WIN. Short activation (5 minutes) of CB1 receptors by bath application of WIN slightly increased in NP bouton density (Fig. 3F,G). Although the increase appeared more robust compared to the 2-AG-induced effect, the effect was too small to reach statistical significance. Similar to 2-AG, the average NP presence appeared slightly increased (Fig. 3H), but we could not detect any changes in specific NP boutons subgroups (Fig. 3I). Together these observations indicate that short CB1 receptor activation by 2-AG or WIN leads to only a small (if any) increase in NP bouton density in GFP-labeled inhibitory axons.

Endocannabinoids are produced on demand in postsynaptic neurons (Alger and Kim, 2011; Hashimotodani et al., 2013; Piomelli, 2014), but an ambient level of endocannabinoids is always present, even in slices (Lee et al., 2015; Lenkey et al., 2015; Szabó et al., 2014). Tonic CB1 receptor activation by endocannabinoids affects mostly perisomatic inhibitory synapses, while dendritic inhibitory synapses are reported to be less sensitive (Lee et al., 2015, 2010). To address if tonic activation of CB1 receptors could have obscured the effects of CB1 receptor activation on inhibitory bouton dynamics in our GFP-labeled axons (which mostly target dendrites (Wierenga et al., 2010)), we applied 5 μM AM251, an antagonist of CB1 receptors. However, AM251 had no effect on NP bouton density (Fig. 3J,K), NP presence or NP bouton subgroups (Fig. 3L,M).

Together our experimental findings indicate that inhibitory bouton dynamics of the GFP-labeled axons are not under strong tonic endocannabinoid control and that short CB1 receptor activation by 2-AG or WIN only slightly increases NP inhibitory bouton density.

### CB1 receptors regulate inhibitory bouton dynamics specifically in CB1R+ axons

The expression of CB1 receptors largely overlaps with the expression pattern of CCK in GABAergic interneurons (Katona et al., 2006, 1999). These interneurons are partially labeled in the GAD65-GFP mice which we use for our experiments (Wierenga et al., 2010). We previously estimated that ∼50% of the GFP-labeled inhibitory axons express CB1 receptors in our slices (Hu et al., 2019), and this may significantly dilute an effect of CB1 receptor activation on bouton dynamics (Fig. 3). We therefore used *post-hoc* immunostaining immediately after two-photon live imaging to distinguish between axons with and without CB1 receptors (CB1R+ and CB1R-axons respectively; Fig. 4A, B). In accordance with previous reports (Dudok et al., 2015; Mikasova et al., 2008), CB1 receptors covered the entire surface of CB1R+ inhibitory axons and individual CB1R+ axons could be easily traced from the CB1 immunostainings (Fig. 4A, B). In addition, there was significant CB1 background staining, presumably reflecting CB1 receptors in pyramidal cells and glia cells (Bonilla-Del Río et al., 2021). CB1R-axons had a higher bouton density and higher bouton turnover compared to CB1R+ axons (Fig. 4C; see below), supporting the notion that CB1R+ and CB1R-GFP-labeled axons belong to separate subtypes of GABAergic cells.

**Figure 4.**
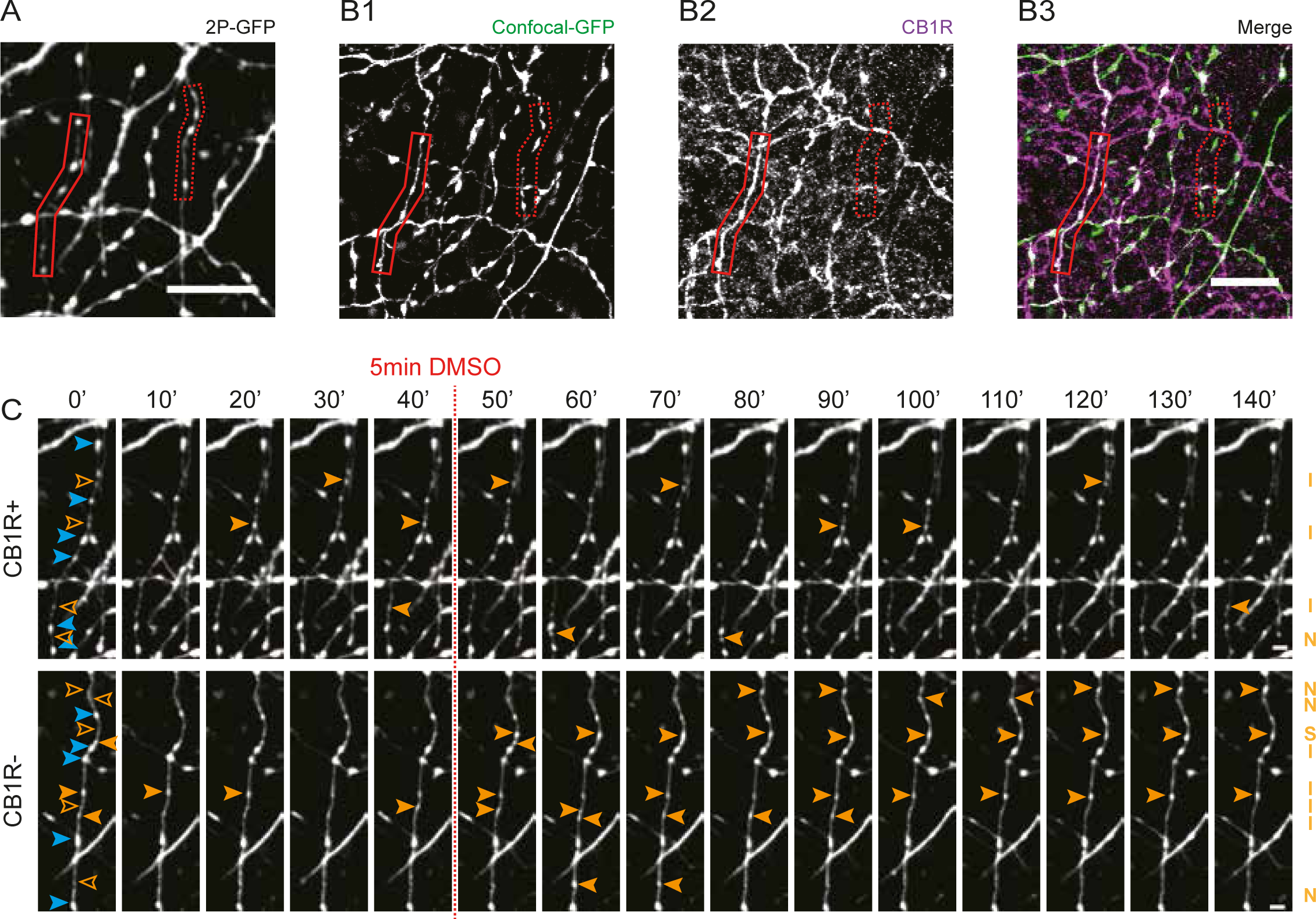
Distinction between CB1R+ and CB1R-axons using post hoc immunohistochemistry. **(A)** Z-projection of representative two-photon image of GFP-labeled inhibitory axons. After two-photon live imaging, the slice was immediately fixated and further processed for immunohistochemistry to assess CB1R expression. **(B)** Confocal images of the same area after post hoc immunohistochemistry, showing the same GFP-labeled axons (B1) as in A (indicated with solid and dashed red boxes). Immunostaining against CB1 receptors (B2) show a clear distinction between CB1R+ axons (solid red box), which express CB1 receptors, which cover the entire axonal surface, and CB1R-axons (dashed red box). Which do not express CB1 receptors. **(C)** Two-photon time lapse imaging of bouton dynamics in the CB1R+ and CB1R-axons indicated in A and B. Arrow heads indicate P (blue) and NP (orange) boutons as in Fig. 3A. Scale bars are 10 µm in A,B and 2 μm in C,D.

We repeated the WIN application experiments, but now separately analyzed CB1R+ and CB1R-axons. In CB1R+ axons the density of NP boutons significantly increased after WIN application (Fig. 5A,B). WIN also increased average NP presence compared to control axons (Fig. 5C). When we analyzed the NP bouton subgroups we found a specific increase in the density of new and stabilizing boutons (Fig. 5D,F), whereas other NP subgroups were unaffected (Fig. 5D-H). New boutons reflect immature synapses, which start to recruit pre- and postsynaptic proteins, while levels of VGAT and gephyrin at stabilizing boutons at the end of the imaging period is comparable to persistent boutons (Frias et al., 2019; Schuemann et al., 2013). In clear contrast, WIN had no effect on bouton density or dynamics in CB1R-axons in the same slices (Fig. 5I-P). These results indicate that axonal CB1 receptors are required for mediating the WIN-induced changes in bouton dynamics in inhibitory axons, and exclude a role for CB1 receptors on other cells. Our results indicate that short activation of axonal CB1 receptors leads to an increase in NP bouton density by specifically promoting the formation and stabilization of inhibitory boutons.

**Figure 5.**
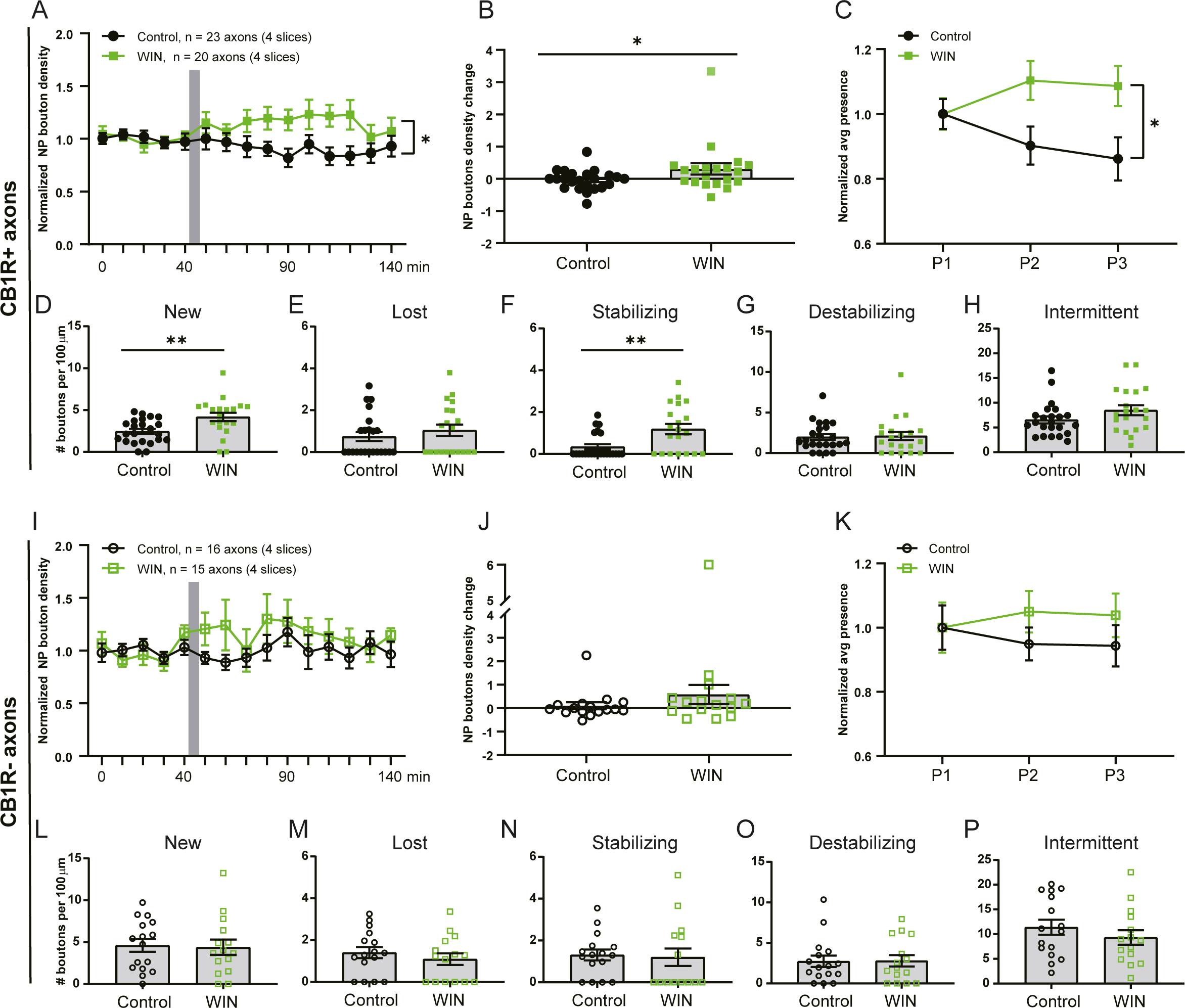
WIN promotes formation and stabilization of inhibitory boutons only in CB1R+ axons. **(A)** Normalized NP bouton density in CB1R+ axons over time in control (black) slices and after WIN (green) application (2w ANOVA, *p* = 0.018; interaction *p =* 0.026). **(B)** Maximum change in NP bouton density in CB1R+ axons in control (black) slices and after WIN (green) application (MW, *p* = 0.047). **(C)** Normalized NP presence in CB1R+ axons over time in control slices and after WIN application (2w ANOVA, *p* = 0.022; interaction *p =* 0.045). **(D-H)** Mean density of NP bouton subgroups in CB1R+ axons in control slices and after WIN application. D, new boutons (MW, *p* = 0.002); E, lost boutons (MW, *p* = 0.39); F, stabilizing boutons (MW, *p* = 0.005); G, destabilizing boutons (MW, *p* = 0.87); H, intermittent boutons (MW, *p* = 0.16). **(I)** Normalized NP bouton density in CB1R-axons over time in control (black) slices and after WIN (green) application (2w ANOVA, *p* = 0.27). **(J)** Maximum change in NP bouton density in CB1R-axons in control (black) slices and after WIN (green) application (MW, *p* = 0.21). **(K)** Normalized NP presence in CB1R-axons over time in control slices and after WIN application (2w ANOVA, *p* = 0.37). **(L-P)** Mean density of NP bouton subgroups in CB1R-axons in control slices and after WIN application. L, new boutons (MW, *p* = 0.77); M, lost boutons (MW, *p* = 0.46); N, stabilizing boutons (MW, *p* = 0.50); O, destabilizing boutons (MW, *p* = 0.99); P, intermittent boutons (MW, *p* = 0.34). Data from 25 CB1R+ and 16 CB1R-axons in 4 control slices and 20 CB1R+ and 15 CB1R-axons in 4 slices with WIN application.

### WIN-induced bouton formation does not require G_i/o_ signaling and neuronal activity

CB1 receptors are G-protein coupled receptors. Endocannabinoid signaling via CB1 receptors typically activates G_i/o_ heterotrimeric proteins, resulting in a reduction of neurotransmitter release at presynaptic terminals (Castillo et al., 2012; Lovinger, 2008). We therefore tested whether WIN-induced bouton formation requires G_i/o_ signaling. We pretreated the slices with pertussis toxin (PTX) (1 μg/ml) for 24 hours to eliminate G_i/o_ signaling (Campbell and Smrcka, 2018; Guo and Ikeda, 2004), and then performed two-photon time-lapse live imaging as before. Axons with and without CB1R were distinguished using *post-hoc* immunostaining (Fig. 6A-C). PTX pretreatment had no major effect on CB1 receptor expression patterns.

**Figure 6.**
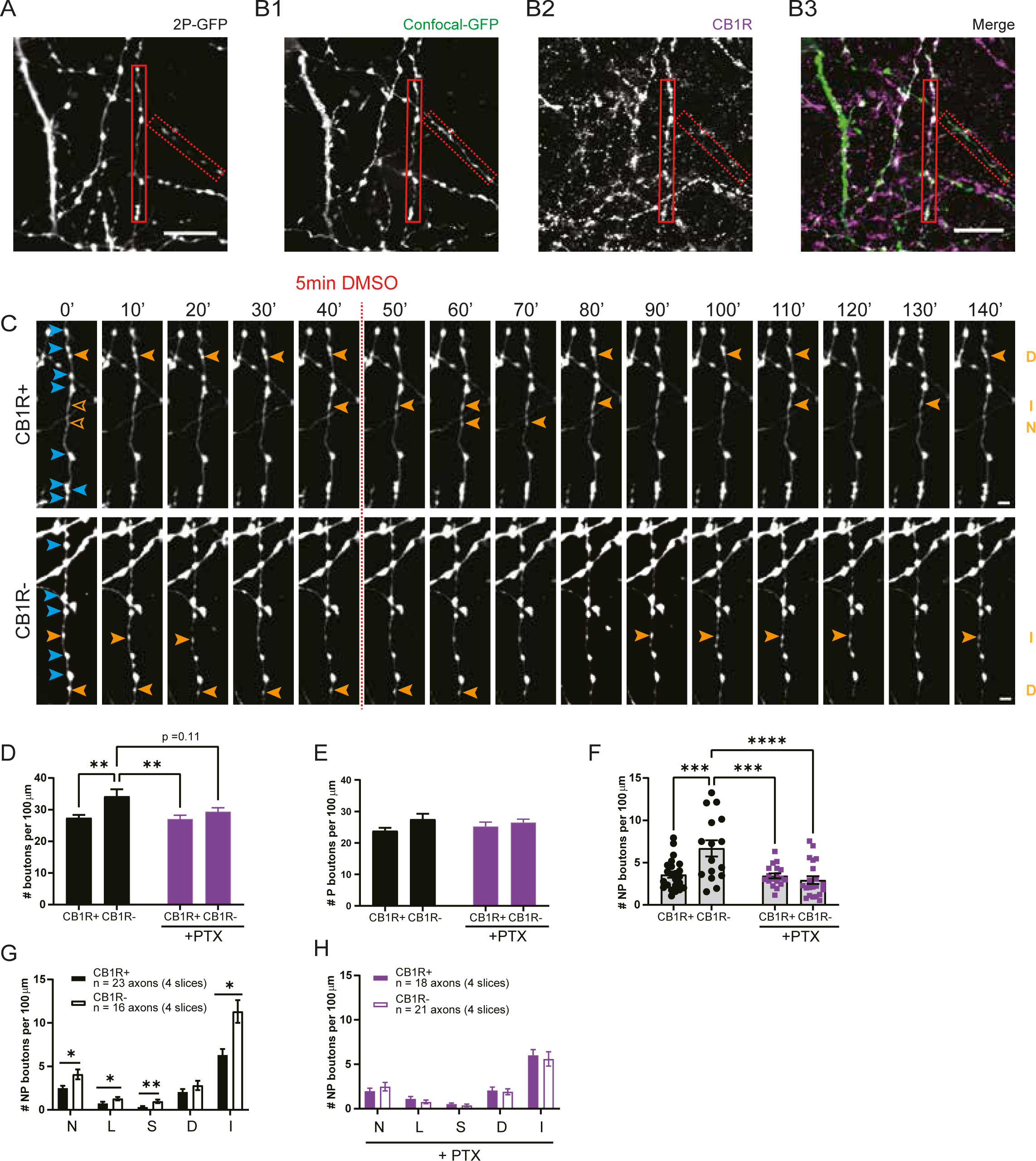
G_i/o_ signaling is an important regulator of inhibitory bouton dynamics. **(A)** Z-projection of representative two-photon image of GFP-labeled inhibitory axons after PTX pretreatment. **(B)** Confocal images of the same area after post hoc immunohistochemistry, showing the same GFP-labeled axons (B1) as in A (solid and dashed red boxes indicate CB1R+ and CB1R-axons). **(C)** Two-photon time lapse imaging of bouton dynamics in the CB1R+ and CB1R-axons indicated in A and B after PTX pretreatment. Arrow heads indicate P (blue) and NP (orange) boutons as in Fig. 3A. **(D)** Average bouton density during baseline in CB1R+ and CB1R-axons in control slices and after PTX pretreatment. Comparisons between CB1R+ and CB1R-axons: *p* = 0.0056 for control, *p* = 0.79 after PTX; between control and PTX: *p* = 0.11 for CB1R-axons, *p* > 0.99 for CB1R+ axons; between CB1R+ (control) and CB1R-(PTX): *p* = 0.86 and between CB1R-(control) and CB1R+ (PTX): *p* = 0.0057 (2w ANOVA Sidak). **(E)** Average density of persistent (P) boutons during baseline in CB1R+ and CB1R-axons in control slices and after PTX pretreatment (p=0.057 for axon type, 2w ANOVA Sidak). **(F)** Average density of non-persistent (NP) boutons during baseline in CB1R+ and CB1R-axons in control slices and after PTX pretreatment. Comparisons between CB1R+ and CB1R axons: *p* = 0.0007 for control, *p* = 0.99 after PTX; between control and PTX: *p* < 0.0001 for CB1R-axons, *p* > 0.99 for CB1R+ axons; between CB1R+ (control) and CB1R-(PTX): *p* = 0.93 and between CB1R-(control) with CB1R+ (PTX): *p* = 0.0008 (2w ANOVA Sidak). **(G)** Mean density of NP bouton subgroups in CB1R+ and CB1R-axons in control slices. N – new boutons (MW, *p* = 0.035); L – lost boutons (MW, *p* = 0.037); S – stabilizing boutons (MW, *p* = 0.002); D – destabilizing boutons (MW, *p* = 0.47); I – intermittent boutons (MW, *p* = 0.010). **(H)** Mean density of NP bouton subgroups in CB1R+ and CB1R-axons after PTX pretreatment (MW, *p* = 0.45 (N); *p* = 0.41 (L); *p* = 0.36 (S); *p* = 0.88 (D); *p* = 0.40(I)). Data from 23 CB1R+ and 16 CB1R-axons in 4 control slices, and 18 CB1R+ and 21 CB1R-xons in 4 PTX-pretreated slices. Scale bars are 10 µm in A,B and 2 μm in C,D.

Under control conditions, CB1R-axons had a higher bouton density compared to CB1R+ axons (Fig. 6D), which was mainly due to a higher density of NP boutons (Fig. 6E,F). The density for all NP bouton subgroups was almost twice as high in CB1R-axons compared to CB1R+ axons (Fig. 6G), showing that overall inhibitory bouton dynamics were more pronounced in CB1R-axons compared to CB1R+ axons. Unexpectedly, we observed that 24 hr pretreatment with PTX affected bouton density. PTX pretreatment specifically downregulated bouton density in CB1R-axons, while bouton density in CB1R+ axons was largely unaffected (Fig. 6D). PTX specifically reduced the density of non-persistent boutons in CB1R-axons (Fig. 6E,F). After PTX pretreatment there was no longer a difference in NP bouton subgroups between CB1R+ and CB1R-inhibitory axons (Fig. 6H). This suggests that under normal conditions CB1R-axons have a higher G_i/o_ protein activity compared to CB1R+ axons in these slices. These data imply that G_i/o_ signaling is an important regulator of inhibitory bouton dynamics.

We then tested whether acute activation of CB1 receptors via WIN can induce changes in inhibitory bouton dynamics in the absence of G_i/o_ signaling. We observed that short activation of CB1 receptors by WIN could still induce the formation of new inhibitory boutons in CB1R+ axons after pretreatment with PTX (Fig. 7A). This indicates that the formation of new inhibitory boutons by CB1 receptor activation is independent of G_i/o_ signaling. However, in the absence of G_i/o_ signaling WIN application no longer promoted bouton stabilization (Fig. 7B; compare with Fig. 5F), suggesting that bouton stabilization requires intact G_i/o_ signaling. As before, other NP bouton subgroups were not affected (Fig. 7C) and WIN application did not affect bouton formation (density of new boutons was 81 ± 23 % of control; MW, p = 0.51) or bouton dynamics (data not shown) in CB1R-axons. These data indicate that short activation of CB1 receptors on inhibitory axons by WIN promotes the formation of new boutons via a G_i/o_-independent signaling pathway.

**Figure 7.**
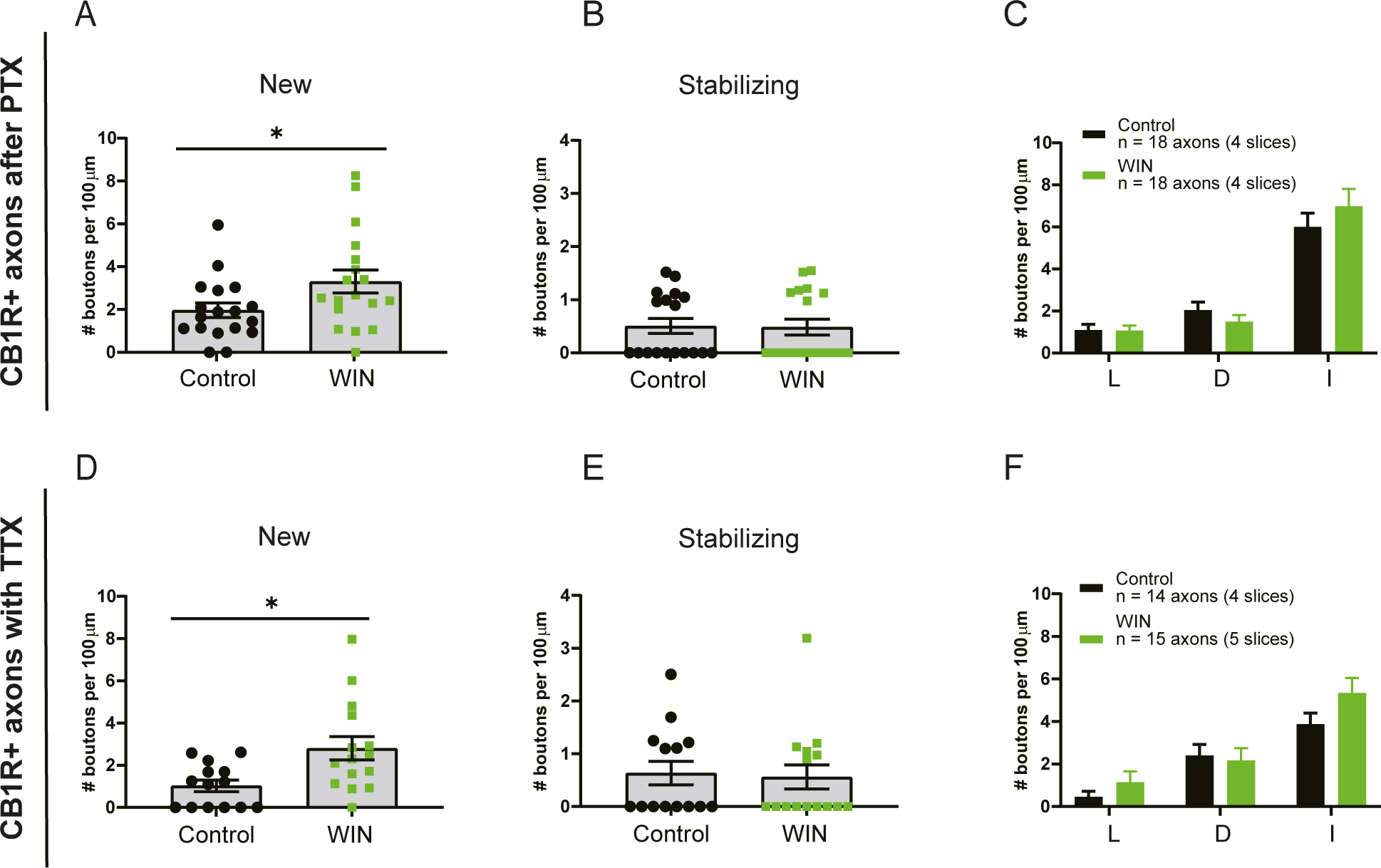
CB1-mediated bouton formation does not require G_i/o_ signaling and is independent of activity. **(A)** Mean density of new boutons in CB1R+ axons after control (black) and WIN (green) application in PTX-pretreated slices (MW, *p* = 0.047). **(B)** Mean density of stabilizing boutons in CB1R+ axons after control and WIN application in PTX-pretreated slices (MW, *p* = 0.93). **(C)** Mean density of other NP bouton subgroups in CB1R+ axons after control and WIN application in PTX-pretreated slices. L – lost boutons (MW, *p* = 0.82); D – destabilizing boutons (MW, *p* = 0.37); I – intermittent boutons (MW, *p* = 0.59). **(D)** Mean density of new boutons in CB1R+ axons after control (black) and WIN (green) application in the presence of TTX (MW, *p* = 0.013). **(E)** Mean density of stabilizing boutons in CB1R+ axons after control and WIN application in the presence of TTX (MW, *p* = 0.61). **(F)** Mean density of other NP bouton subgroups in CB1R+ axons after control and WIN application in the presence of TTX. L – lost boutons (MW, *p* = 0.23); D – destabilizing boutons (MW, *p* = 0.56); I – intermittent boutons (MW, *p* = 0.16). Data in A-C from 18 axons in 4 slices with DMSO (control) application and 18 axons in 4 slices with WIN application. Data in D-F from 14 axons in 4 slices with DMSO (control) application and 15 axons in 5 slices with WIN application.

G_i/o_ protein signaling can hyperpolarize neurons via activation of K^+^ channels (Bacci et al., 2004; Guo and Ikeda, 2004). Blocking ongoing G_i/o_ activity with PTX may therefore enhance neuronal activity in our slices, which may by itself affect inhibitory bouton dynamics. However, as enhancing neuronal activity is expected to promote overall inhibitory bouton turnover (Frias et al., 2019; Schuemann et al., 2013), this does not appear in line with the observed decrease in inhibitory bouton dynamics in CB1R-axons after PTX. To address if WIN-induced inhibitory bouton formation is affected by activity, we blocked network activity with TTX to reduce overall bouton dynamics (Frias et al., 2019; Schuemann et al., 2013). We observed that in the presence of TTX, brief activation of CB1 receptors with WIN still induced the specific increase in the density of new boutons (Fig. 7D). However, WIN did no longer induce a change in the density of stabilizing boutons (Fig. 7E), consistent with our earlier finding that inhibitory bouton stabilization requires activity (Frias et al., 2019). Other NP bouton subgroups were not affected (Fig. 7F) and WIN application did not significantly affect bouton formation (179 ± 216 % of control; MW, p = 0.11) or other bouton dynamics (data not shown) in CB1R-axons. Together these data demonstrate that CB1 receptor-mediated inhibitory bouton formation does not require G_i/o_ protein signaling and is independent of neuronal activity.

### Acute elevation of cAMP levels promotes inhibitory bouton formation

Besides the typical downstream signaling pathway via G_i/o_ proteins, CB1R activation can trigger several other signaling pathways, including via G_12/13_ (Roland et al., 2014), G_q_ (Lauckner et al., 2005) and G_s_ proteins (Finlay et al., 2017; Glass and Felder, 1997). Intriguingly, a novel form of CB1 receptor-mediated synaptic potentiation was recently reported, which was shown to depend on presynaptic PKA activity (Cui et al., 2016; Wang et al., 2017). This raises the attention to CB1 receptor-mediated G_s_ signaling, as G_s_ protein signaling enhances PKA activity via stimulation of cAMP production (Antoni, 2012; Taylor et al., 2013). We therefore tested if inhibitory bouton dynamics were affected when we directly elevated cAMP levels via activation of adenylyl cyclase (AC) by 25 μM forskolin (5 minutes) (Fig. 8A). We observed that brief application of forskolin induced the formation of new inhibitory boutons (Fig. 8B), while other NP subgroups were not affected (Fig. 8C,D). This suggests that the inhibitory bouton formation that we observed after CB1 receptor activation may be mediated by G_s_ signaling. The increase in inhibitory bouton formation after forskolin application appeared much stronger compared to WIN application (compare Fig. 8B to Fig. 3I), suggesting that most, if not all, GFP-labeled inhibitory axons responded to forskolin. These data show that the formation of new inhibitory boutons is promoted by increasing intracellular cAMP levels via AC stimulation.

**Figure 8.**
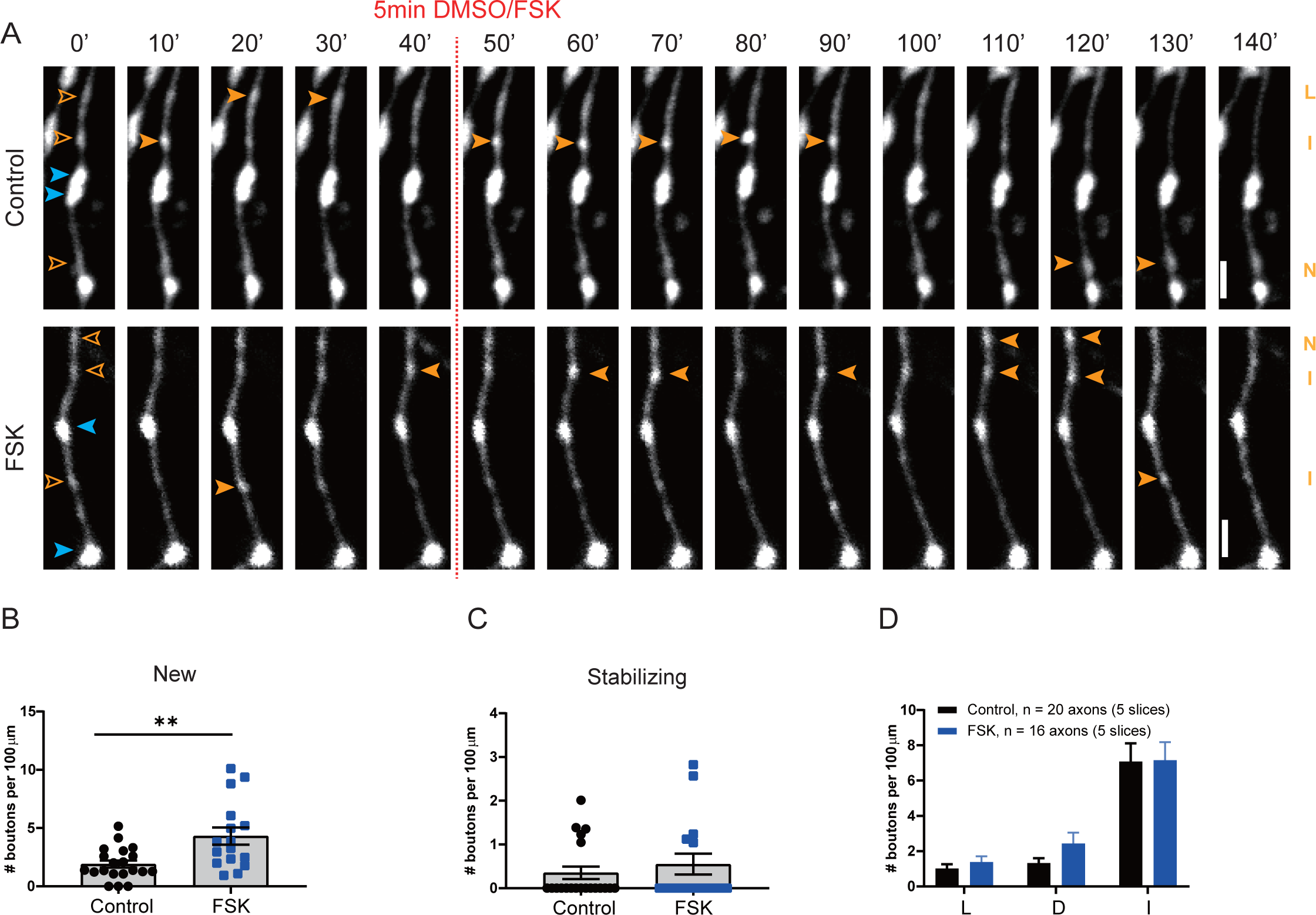
Inhibitory bouton formation is promoted by increasing intracellular cAMP levels with forskolin. **(A)** Representative two-photon time lapse images of bouton dynamics in GFP-labeled axons after control or forskolin application. Arrow heads indicate P (blue) and NP (orange) boutons as in Fig. 3A. Scale bar is 2 µm. **(B)** Mean density of new boutons in control (black) slices and after forskolin (blue) application (MW, *p* = 0.007). **(C)** Mean density of stabilizing boutons in control slices and after forskolin application (MW, *p* = 0.67). **(D)** Mean density of other subgroup of NP boutons in control slices and after forskolin application. L – lost boutons (MW, *p* = 0.46); D – destabilizing boutons (MW, *p* = 0.37); I – intermittent boutons (MW, *p* = 0.81). Scale bars are 2 μm.

### G_s_ signaling in inhibitory axons promotes inhibitory bouton formation

Bath application of forskolin strongly increases neuronal activity (Gekel and Neher, 2008; Mitoma and Konishi, 1996) and will raise cAMP levels in all cells in the slice. The observed specific increase in new inhibitory bouton formation after forskolin, without affecting overall bouton dynamics, is therefore quite remarkable. However, we cannot conclude that the observed increase in inhibitory bouton formation is a direct effect of elevated cAMP levels in the inhibitory axons. We made use of DREADDs (Designer Receptors Exclusively Activated by Designer Drugs) (Roth, 2016; Urban and Roth, 2015) to achieve cell-specific manipulation of presynaptic cAMP levels. G_s_-DREADDs allow the direct activation of the G_s_-protein signaling pathway using the specific ligand CNO. To achieve sparse expression restricted to inhibitory neurons we infected hippocampal slices from VGAT-Cre mice with Cre-dependent AAVs. We used two AAVs: one containing a HA-tagged G_s_-DREADD construct and one containing GFP (Fig. 9A; see methods for details). Infections with these two AAVs resulted in sparse GFP-labeling of inhibitory cells and their axons, which partially overlapped with G_s_-HA expression (Fig. 9B). P*ost-hoc* immunostaining allowed us to identify GFP-labeled axons with and without G_s_-HA (HA+ and HA-axons) in the same slice (Fig. 9C,D). We performed two-photon microscopy to monitor bouton dynamics in GFP-expressing HA+ and HA-inhibitory axons (Fig. 9E). Bouton dynamics in VGAT-Cre slices were in line with previous data (Frias et al., 2019), indicating that the AAV infections did not alter overall bouton dynamics in inhibitory axons. After a 40-minute baseline period, G_s_-DREADDs were activated via bath application of CNO ligand. We found that CNO activation strongly increased the density of new boutons in G_s_-HA positive axons compared to HA-axons (Fig. 9F). Other NP bouton subgroups were not affected, although the density of stabilizing boutons appeared to be somewhat increased (Fig. 9G,H). These data show that specific activation of G_s_ signaling in inhibitory axons mimics the WIN-induced inhibitory bouton formation.

**Figure 9.**
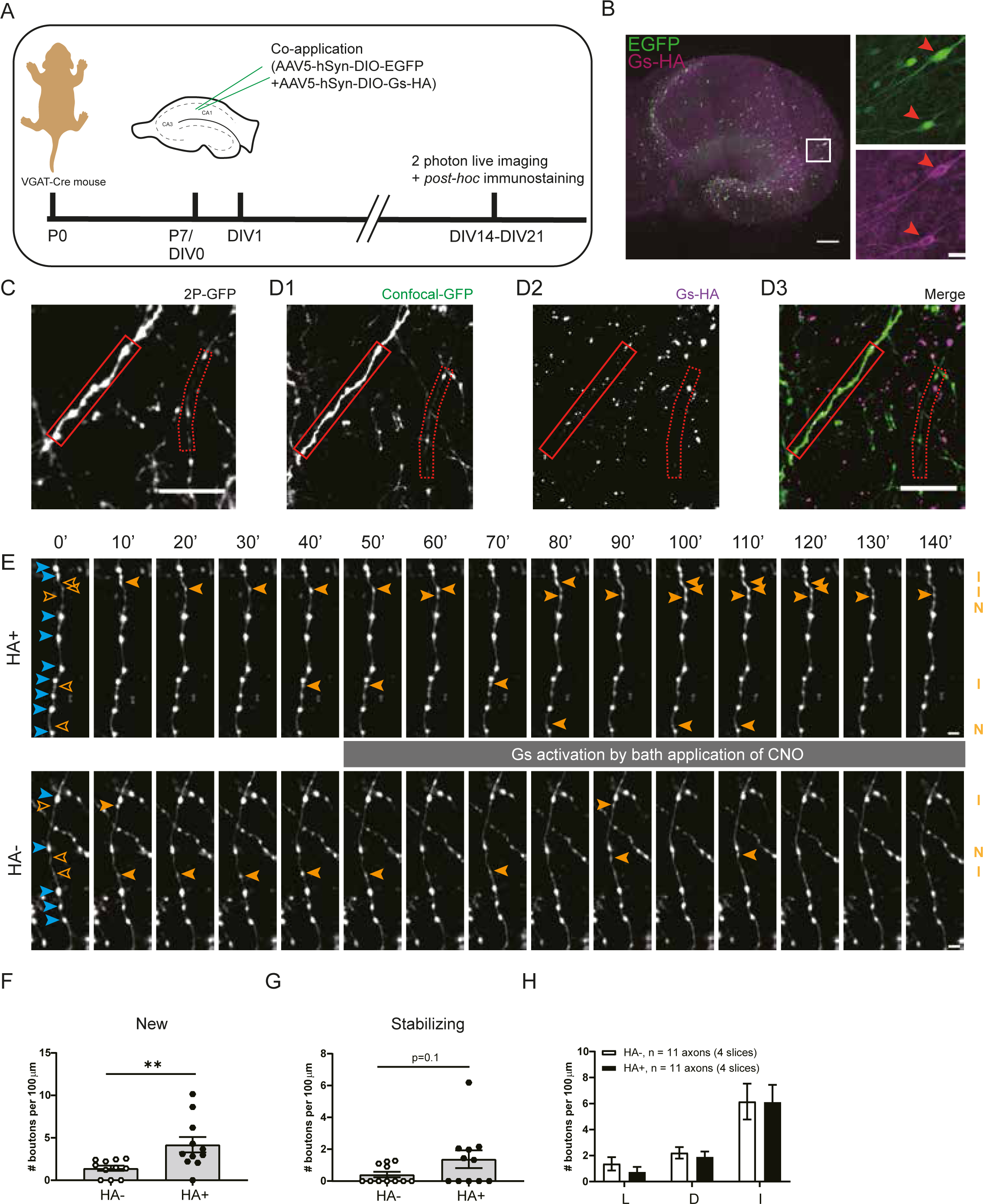
Specific activation of Gs at inhibitory axons induce new bouton formation. **(A)** Experimental design. Hippocampal slice cultures are prepared from P7 VGAT-Cre mouse pups. At DIV1 (days *in vitro*), AAV5-hSyn-DIO-EGFP and AAV5-hSyn-DIO-Gs-HA viruses are applied to the VGAT-Cre slice cultures. After 2-3 weeks (DIV 14-21) slices were used for two-photon live imaging and *post hoc* immunostaining to reveal Gs-HA expression. **(B)** Representative example of VGAT-Cre slice culture at DIV20 showing sparse expression of GFP and Gs-HA in GABAergic cells. Right images (zoom from white box) show Gs-HA and EGFP co-expression in a subset of neurons (red arrow heads). **(C)** Z-projection of representative two-photon image of GFP-labeled inhibitory axons in VGAT-Cre slice. **(D)** Confocal images of the same area in C after post hoc immunohistochemistry against HA, showing the same GFP-labeled axons as in A (solid and dashed red boxes indicate HA+ and HA-axons). **(E)** Two-photon time lapse imaging of bouton dynamics in the HA+ and HA-axons indicated in C and D. Gs-DREADDs were activated by bath application of 10 µM CNO after the 40 minutes baseline period. Arrow heads indicate P (blue) and NP (orange) boutons as in Fig. 3A. **(F)** Mean density of new boutons at HA+ and HA-axons in response to Gs-DREADD activation (MW, *p* = 0.003). **(G)** Mean density of stabilizing boutons at HA+ and HA-axons in response to Gs-DREADD activation (MW, *p* = 0.10). **(H)** Mean density of other subgroup of NP boutons at HA+ and HA-axons in response to Gs-DREADD activation. L – lost boutons (MW, *p* = 0.30); D – destabilizing boutons (MW, *p* = 0.44); I – intermittent boutons (MW, *p* = 0.85) Data from 11 HA+ and 11 HA-axons in 4 slices. Scale bars are 200 μm (overview) and 20 μm (zoom) in B, 10 μm in C,D and 2 μm in E.

Together, our results indicate that inhibitory bouton formation after brief CB1 receptor activation does not require G_i/o_-signaling, and that it is mimicked by activation of G_s_ signaling in inhibitory axons. This suggests that CB1 receptors on inhibitory axons couple to G_s_ proteins rather than the conventional G_i/o_ effectors to trigger inhibitory bouton formation.

## Discussion

Here we examined the signaling pathway underlying the CB1 receptor-mediated formation of new inhibitory synapses. We made several important observations. First of all, repeated CB1 activation led to an increase in mIPSC frequency and an increase in the density of presynaptic VGAT clusters, which were not associated with postsynaptic gephyrin. Inhibitory synapses which do not contain gephyrin are immature and show reduced transmission (Danglot et al., 2003; Nguyen et al., 2016; Niwa et al., 2012; Patrizi et al., 2008; Yu et al., 2007). Our observations are in line with the notion that pre- and postsynaptic signaling pathways during synapse formation are largely independent (Jiang et al., 2021; Wierenga, 2017) and suggest that CB1 receptors act purely presynaptically. Second, brief activation of CB1 receptors specifically triggered the formation of inhibitory synapses in CB1R+ axons. This indicates that formation of inhibitory synapses is mediated by axonal CB1 receptors and excludes a prominent role for CB1 receptors in astrocytes or postsynaptic neurons. Third, bouton turnover in inhibitory axons was strongly reduced when G_i/o_ protein signaling was blocked by PTX pretreatment. This suggests that modulation of axonal cAMP levels is an important regulator of bouton turnover in inhibitory axons. Fourth, CB1 receptor-mediated inhibitory bouton growth was independent of ongoing G_i/o_ signaling and activity, suggesting that signaling pathways downstream of axonal CB1 receptors differ from presynaptic CB1 receptors. Finally, inhibitory synapse formation was induced in response to an increase in cAMP after forskolin application, and when G_s_ signaling was activated via G_s_-DREADDs, which were expressed exclusively in inhibitory neurons. These findings revealed that an increase in cAMP is the key second messenger signal for inhibitory bouton formation and suggest that axonal CB1 receptors trigger inhibitory bouton formation via G_s_ instead of G_i/o_ protein signaling.

Our present study has limitations that are important to mention here. First of all, we use transgenic mice in which several inhibitory neuron subtypes are labeled with GFP (Wierenga et al., 2010). This unspecific labeling diluted and hampered the detection of axon-specific effects (Fig. 2). However, we used it to our advantage by performing posthoc immunostaining to distinguish between different inhibitory axon types. This allowed comparison between CB1R+ and CB1R-, or HA+ and HA-axons in the same slice and avoided comparison between slices from different GFP-labeled mouse lines. Another limitation of our study is that we have used bath application of CB1 agonist WIN to trigger inhibitory bouton formation. Under physiological conditions, endocannabinoid signals are likely transient and highly localized (Hashimotodani et al., 2007; Hu et al., 2019; Monday and Castillo, 2017), providing spatial and temporal control over inhibitory synapse formation. We elevated cAMP levels to trigger inhibitory bouton formation by bath application of forskolin or by activation of G_s_-

DREADDs in inhibitory cells. While this allowed separation of formation and stabilization of inhibitory boutons, it likely abolished spatial modulations. Axons contain several phosphodiesterases, which rapidly degrade cAMP and provide spatiotemporal compartmentalization of cAMP signaling (Argyrousi et al., 2020; Baillie, 2009). Pretreatment with PTX will disturb these cAMP modulations and this strongly reduced inhibitory bouton dynamics and abolished the difference between CB1R+ and CB1R-axons (Fig. 6L). This indicates that CB1R-axons have higher G_i/o_ baseline activity compared to CB1R+ axons and suggests that cAMP modulation is an important factor regulating inhibitory bouton dynamics. Future research should further assess the relationship between cAMP signaling and inhibitory bouton turnover.

Synapse formation is a multistep process, with each step regulated by specific signaling pathways (Jiang et al., 2021; Wierenga, 2017). Our detailed two-photon analysis allows dissecting these steps and addressing the involved signaling pathways. Inhibitory synapse formation starts with the growth of a new bouton at an axonal location where the inhibitory axon is in close proximity to a dendrite (Dobie and Craig, 2011; Hu et al., 2019; Villa et al., 2016; Wierenga et al., 2008). Our data indicate that axonal CB1 receptors can trigger bouton formation, which does not require neuronal activity. We observed that CB1 receptor-mediated inhibitory bouton formation was not affected in the presence of TTX (Fig. 7D). In addition, we observed that forskolin, which strongly raises neuronal activity (data not shown), did not affect overall bouton turnover (Fig. 8D). This was unexpected given our previous observations that inhibitory bouton turnover is enhanced by neuronal activity (Frias et al., 2019; Schuemann et al., 2013). On the other hand, we observed that blocking G_i/o_ signaling strongly affected bouton turnover (Fig. 6F,H), which appeared independent of activity. These data suggest that axonal cAMP is the primary second messenger affecting inhibitory bouton formation, which is indirectly modulated by activity, possibly via changes in neuromodulatory signals.

Our data indicate that axonal CB1 receptors can directly trigger bouton formation via an increase in cAMP, while subsequent bouton stabilization and postsynaptic assembly requires additional signaling. WIN-induced bouton stabilization was prevented when G_i/o_ signaling was blocked by PTX (Fig. 7B), and bouton stabilization was not altered by increasing cAMP levels with forskolin (Fig. 8C), although it may be facilitated with longer elevations (Fig. 9H). These data suggest that after the initial formation, CB1 receptors may also promote bouton stabilization via a more indirect pathway. We previously showed that bouton stabilization requires neuronal activity and involves local actin remodeling via a reduction in ROCK activity (Frias et al., 2019). Interactions between CB1 receptor signaling and ROCK activity (Berghuis et al., 2007) and actin remodeling (Njoo et al., 2015; Zhou et al., 2019) have been reported, but future research should further clarify the precise nature of these interactions.

CB1 receptors are highly versatile and are involved in many neuronal processes via multiple downstream pathways, including axon guidance and synaptic plasticity (Araque et al., 2017; Berghuis et al., 2007; Monday and Castillo, 2017; Njoo et al., 2015; Roland et al., 2014). There are multiple factors, including interacting proteins (Guggenhuber et al., 2016), which determine which downstream signaling pathway is activated after CB1 receptor activation (Flores-Otero et al., 2014; Nogueras-Ortiz and Yudowski, 2016) and this functional selectivity of CB1 receptors may have important clinical relevance (Ibsen et al., 2017; Laprairie et al., 2017; Sholler et al., 2020). It was recently reported that the duration of CB1 receptor activation determines the direction of plasticity at corticostriatal synapses with brief activation inducing LTP, while prolonged activation induces LTD (Cui et al., 2016, 2015). Our data suggest that brief activation of axonal CB1 receptors promotes the formation of new inhibitory boutons via G_s_-mediated elevation of cAMP levels, but we have not extensively tested longer activations or different ligand concentrations. It is possible that the subcellular location of CB1 receptors affects downstream signaling pathway: CB1 receptors at presynaptic terminals couple to G_i/o_ to affect GABA release (Guo and Ikeda, 2004; Lee et al., 2015), while CB1 receptors in the axonal shaft of the same inhibitory axons may couple to G_s_ proteins. Even though CB1 receptors prefer coupling to G_i_-proteins, they can switch to G_s_ when G_i_-proteins are not available or already occupied (Caballero-Florán et al., 2016; Eldeeb et al., 2016; Finlay et al., 2017; Glass and Felder, 1997). This may suggest that G_i_ proteins are only available at presynaptic terminals, while G_s_ protein coupling could be dominant in axons.

Our experiments do not address which effectors are downstream of cAMP to trigger inhibitory bouton formation. The most prominent effector of cAMP is protein kinase A (PKA). Presynaptic PKA activity is involved in CB1-mediated synaptic plasticity (Chevaleyre et al., 2007; Cui et al., 2016) and also is therefore a strong candidate for regulating presynaptic bouton formation. PKA may for instance alter local clustering and inter-bouton exchange of synaptic vesicles (Chenouard et al., 2020; Patzke et al., 2019). PKA resides close to the plasma membrane and preferably phosphorylates membrane proteins in its close proximity (Tillo et al., 2017). However, potential PKA targets to mediate inhibitory bouton formation remain yet to be identified. In addition, there are important PKA-independent pathways downstream of cAMP signaling, most importantly via Epac2 (Kawasaki et al., 1998). Epac2 activity can strongly increase synaptic transmission (Fernandes et al., 2015; Gekel and Neher, 2008), yet a role in synapse formation has not been reported. Interestingly, Epac2 was recently found to be downstream of G_s_-coupled β adrenergic receptors to mediate presynaptic LTP at parallel fiber synapses to Purkinje cells (Martín et al., 2020). cAMP signaling via PKA, Epac2 or Rho GTPases may affect the axonal cytoskeleton. Actin is important in the formation, stabilization and maintenance of presynaptic terminals (Bednarek and Caroni, 2011; Chenouard et al., 2020; Chia et al., 2014, 2013; Frias et al., 2019; Pielage et al., 2011) and cAMP fluctuations may drive local modifications in the actin cytoskeleton (Bernier et al., 2019) underlying structural presynaptic changes.

Our findings suggest that axonal CB1 receptors serve an important role in local, on demand synapse formation. Our observation that inhibitory bouton formation was more prominent after cAMP elevation than after WIN application (compare Fig. 8B and 9F to 3I) suggests that axonal cAMP signaling is an important second messenger signal mediating bouton formation not only in CB1R+, but perhaps in all, inhibitory axons. Intriguingly, our observations are reminiscent of cAMP-mediated bouton formation in zebrafish (Yoshida and Mishina, 2005), Aplysia (Bailey and Kandel, 1993; Nazif et al., 1991; Upreti et al., 2019) and Drosophila axons (Koon et al., 2011; Maiellaro et al., 2016; Zhong et al., 1992). This raises the possibility that axonal cAMP signaling is a universal second messenger system for regulating structural plasticity in axons. Activation of CB1 receptors via dendritic endocannabinoid signaling (Hu et al., 2019) then represents one specific way to trigger cAMP-mediated bouton formation in CB1R+ axons in response to strong excitatory synaptic activity. Other axons may employ different axonal receptors to mediate bouton formation. Indeed, GABAergic interneurons express many different G-proteins (Cox et al., 2008; Helboe et al., 2015; Puighermanal et al., 2017), which often provide neuromodulatory context signals from other brain areas (Hattori et al., 2017). Our findings raise the intriguing possibility that neuromodulatory receptors on the axonal surface provide the opportunity to build a new inhibitory bouton on demand, triggered by axon-specific and context-dependent signaling.

## Materials and Methods

### Animals

All animal experiments were performed in compliance with the guidelines for the welfare of experimental animals issued by the Federal Government of The Netherlands. All animal experiments were approved by the Animal Ethical Review Committee (DEC) of Utrecht University.

### Mouse hippocampal slice culture

Organotypic mouse hippocampal slices were acquired from female and male GAD65-GFP mice at 6-7 days after birth. In these mice, ∼20% interneurons are labelled by GFP from early embryonic developmental stage into adulthood (López-Bendito et al., 2004). Most GFP-labelled interneurons target dendrites of CA1 pyramidal cells and express VIP or reelin, while parvalbumin and somatostatin-positive neurons are not labelled (Wierenga et al., 2010). Slice culture preparation details are described previously (Frias et al., 2019; Hu et al., 2019). Mice were sacrificed and the isolated hippocampus was placed in ice-cold HEPES-GBSS (containing 1.5 mM CaCl_2_·2H_2_O, 0.2 mM KH_2_PO_4_, 0.3 mM MgSO_4_·7H_2_O, 5 mM KCl, 1 mM MgCl_2_·6H_2_O, 137 mM NaCl, 0.85 mM Na_2_HPO_4_ and 12.5 mM HEPES) supplemented with 12.5 mM HEPES, 25 mM glucose and 1 mM kynurenic acid (pH set around 7.2, osmolarity set around 320 mOsm, sterile filtered). Slices were vertically chopped along the long axis of hippocampus at thickness of 400 µm. They were then quickly washed with culturing medium (consisting of 48% MEM, 25% HBSS, 25% horse serum, 30 mM glucose and 12.5 mM HEPES, pH set at 7.3-7.4 and osmolarity set at 325 mOsm), and transferred to Millicell cell culture inserts (Milipore) in 6-well plates. Slices were cultured in an incubator (35 °C, 5% CO_2_) until use. Culturing medium was completely replaced twice a week. Slices were used after 2 to 3 weeks *in vitro*, when the circuitry is relatively mature and stable (De Simoni et al., 2003).

### Pharmacological treatments

The following drugs were used: 20 µM WIN 55212-2 (WIN; Tocris Bioscience), 100 µM 2-AG (Tocris Bioscience), 25 µM forskolin (Abcam), Pertussis toxin 1 µg/ml (Tocris Bioscience), 10 µM CNO (Tocris Bioscience), 5 µM AM251 (Tocris Bioscience). For acute treatments, ACSF containing the drug or 0.1% DMSO vehicle was bath applied for 5 minutes. AM251 and CNO were bath applied after a baseline period (5 time points) and continued until the end of the experiment. Pertussis toxin was added to the slice culture medium and a small drop was placed on top of the slice 24 hours before the start of the experiment. Treated slices were kept in the incubator.

CB1 receptor activation can result in different downstream signaling pathways, which depend on ligand concentration and duration (Cui et al., 2016, 2015). We used a relatively high concentration of WIN (20 μM) to aim for strong activation of CB1 receptors. We used short applications to mimic CB1 receptor activation under physiological conditions (Hu et al., 2019) and to avoid the induction of synaptic depression (Monday et al., 2020).

For repeated treatment of 2-AG, normal culturing medium was replaced by medium containing 100 µM 2-AG or 0.1% DMSO for 20 minutes. This was repeated 3 times with 2 hours intervals. At the start of each medium replacement, a small drop was placed on top of the slices to facilitate exchange. A treatment duration of 20 minutes (rather than 5) was chosen to ensure penetration in the entire slice as solution exchange may be slower in the incubator compared to the microscope bath. After the last treatment, the medium was replaced 3 times with fresh medium to ensure wash out. During and after the treatment slices were kept in the incubator and experiments (immunocytochemistry or electrophysiology) were performed 24 hours after the start of the first treatment.

### Electrophysiology recording and analysis

Slices were transferred to a recording chamber which was continuously perfused with carbogenated artificial cerebrospinal fluid (ACSF; containing 126 mM NaCl, 3 mM KCl, 2.5 mM CaCl_2_, 1.3 mM MgCl_2_, 1.25 mM NaH_2_PO_4_, 26 mM NaHCO_3_, 20 mM glucose and 1 mM Trolox) at 32 °C. Whole-cell patch clamp measurements were recorded with a MultiClamp 700B amplifier (Molecular Devices) and stored using pClamp 10 software. Recordings were filtered with a 3 kHz Bessel filter. Thick-walled borosilicate pipettes of 4–6 MΩ were filled with pipette solution containing (in mM): 70 K-gluconate, 70 KCl, 0.5 EGTA, 10 HEPES, 4 MgATP, 0.4 NaGTP, and 4 Na_2_-phosphocreatine. Cells were discarded if series resistance was above 35 MΩ or if the resting membrane potential exceeded -50 mV. Recordings were excluded when the series resistance after the recording deviated more than 30% from its original value. To isolate miniature inhibitory postsynaptic currents (mIPSCs) TTX, AP5 and DNQX were added to the ACSF. The mIPSCs were analyzed in pClamp and Matlab with homemade scripts (Ruiter et al., 2020). Rise time of mIPSCs were determined as the time between10% and 90% of the peak value. The distribution of the rise times of mIPSCs recorded in control conditions (generated from 150 mIPSCs per cell) were fitted with two Gaussians and their crossing point determined the separation between fast and slow mIPSCs (Ruiter et al., 2020). A double Gaussian fit for the rise time distribution in 2-AG conditions gave a similar separation value (control: 0.9 ms; 2-AG: 1.1 ms) and we verified that our conclusions did not change by taking the 2-AG separation value.

### Two-photon time lapse imaging

Time-lapse two-photon imaging was performed in carbogenated, continuously perfused ACSF at 32 °C. Slices were transferred in a 3 cm dish containing ACSF. Two-photon imaging was performed on a customized two-photon laser scanning microscope (Femto2D, Femtonics, Budapest, Hungary) with a Ti-Sapphire femtosecond pulsed laser (MaiTai HP, Spectra-Physics) with a 60x water immersion objective (Nikon NIR Apochromat, NA = 1.0). A 4x objective (Nikon Plan Apochromat) was used to determine the location of the dendritic layer of the CA1 region. GFP was excited at 910 nm to visualize GFP-labelled axons. 3D image stacks were acquired at a size of 93.5 µm x 93.5 µm (1124 x 1124 pixels) with 50-63 z-steps (0.5 µm step size). Acquisition time per image stack was ∼7 minutes. We acquired image stacks every 10 minutes, with a total of 15 time points (140 minutes). After a baseline of 5 time points, drugs were bath applied.

For slices in which we performed post-hoc immunostaining, an overview of the imaging region was made after the last time point (203 µm x 203 µm, ∼50 z-steps of µm step size), and a line scar was made using high intensity laser power at 910 nm at the edge of the zoomed out imaging area to facilitate alignment with post-hoc confocal microscopy.

### Two-photon image analysis

Individual axons with at least 50 μm length were traced using the CellCounter plugin imbedded in Fiji for all time points (TPs). Individual boutons were identified with custom-built semi-automatic Matlab software, as described previously (Frias et al., 2019). In short, a 3D axonal intensity profile was reconstructed at each TP and individual boutons were selected based on a local threshold (0.5 standard deviation above mean axon intensity). Only boutons containing at least 5 pixels above threshold were included. All boutons at all time points were visually inspected and manual corrections were made if deemed necessary.

Persistent (P) boutons were defined as boutons which were present at all TPs. Non-persistent (NP) boutons were absent at one or more TPs. Boutons which were present for only one time point were considered transport events and were excluded (Frias et al., 2019; Schuemann et al., 2013). Based on their presence or absence during baseline (first 5 TP) and after treatment, NP boutons were further classified into five subgroups (Frias et al., 2019; Ruiter et al., 2021): new boutons (only present after baseline), lost boutons (only present during baseline), stabilizing boutons (non-persistent during baseline, persistent after treatment), destabilizing boutons (persistent during baseline, non-persistent after treatment), intermittent boutons (non-persistent during baseline and after treatment).

Bouton density was calculated per axon as the average number of boutons at each TP divided by the 3D axon length. NP bouton density was determined for each TP as the number of NP boutons that were present divided by the 3D axon length. NP bouton densities were normalized to the average baseline value (first 5 TP) to allow comparison between axons. The maximum change in NP bouton density change was calculated as the maximum NP bouton density (average over 3TPs) divided by the baseline NP bouton density (average over TP2-4). NP presence was determined as the fraction of NP boutons that were present at each time point and these values were averaged for the first, second and third period of 5 TPs each. Changes in NP presence reflect changes in the density of NP bouton subgroups, as well as in bouton duration. However, differences in bouton duration (% of TPs present) of NP bouton subgroups were never observed in any of the conditions and we therefore only report NP bouton densities.

### Immunocytochemistry and confocal microscopy

Fixation of the slices was performed in 4% paraformaldehyde (PFA) for 30 minutes at room temperature covered by aluminum foil. Following washing with phosphate-buffered saline (PBS; 3x 10 minutes), slices were permeabilized with 0.5% Triton X-100 (15 minutes), followed by PBS washing (3x 5 minutes), and 1 hour incubation in blocking solution (0.2% Triton X-100 and 10% goat serum). Application of primary antibodies in blocking solution was performed overnight at 4 °C. After PBS washing (3x 15 minutes), secondary antibodies were applied for 4 hours. After PBS washing (2x 15 minutes), slices were mounted in Vectashield solution.

We used the following primary antibodies: rabbit α-VGAT (Synaptic Systems, 131 003; 1:1000), mouse α-gephyrin (Synaptic Systems, 147 011; 1:1000), mouse α-CB1R (Synaptic Systems, 258 011; 1:1000), rat α-HA (Roche, 11 867 423 001; 1:500), and the following secondary antibodies: anti-mouse Alex647 (Life Technologies, A21241, 1:500) and anti-rabbit Alex405 (Life Technologies, A31556, 1:250) for VGAT and gephyrin staining, anti-mouse Alexa647 (Life Technologies, A21236, 1:500) and anti-mouse Alexa568 (Life Technologies, A11031, 1:500) for CB1R staining, and anti-rat Alexa568 (Life Technologies, A11077, 1:500) for HA staining.

Confocal imaging was performed using a Zeiss LSM-700 microscope system with a 63x oil-immersion objective. A 20x objective was used to find back the two-photon imaging area based on the line scar. Image size was 101.3 µm x 101.3 µm (1024 x 1024 pixels) with 0.3 µm z steps for synapse quantification, and up to 203 µm x 203 µm for post-hoc axon identification.

Confocal images were analyzed in Fiji and corresponding axons in the confocal and two-photon images were identified using the line scar as a guide. Expression of CB1R or HA was determined by visual inspection. In some cases, the image was mirrored to confirm or reject positive staining. Negative axons were always chosen close to positive axons in the same imaging area, assuring that the absence of CB1R or HA expression was not due to low immunostaining quality. In addition, we verified that CB1R expression or staining levels did not affect our conclusion as we found the same results when we split CB1R+ axons in two separate groups with high and low CB1R levels. Per slice, 2-6 axons per group were included in the analysis.

For synapse quantification images were analyzed in Fiji using a custom macro (Ruiter et al., 2020). An average projection image was made from 5 z-planes, images were median filtered (1 pixel radius) and individual puncta were identified using watershed segmentation. VGAT and gephyrin puncta were analyzed separately and overlap was determined afterwards. Four independent experiments were performed with 1 or 2 image areas per slice. To compare between treatments, data were normalized per experiment.

### VGAT-Cre slice preparation and AAV virus injection

Hippocampal slice cultures were prepared as described above from VGAT-Cre mice (JAX stock #028862) at 6-7 days after birth. Floxed AAV5 viruses (DIO-EGFP, #v115-1, DIO-Gs-HA, #v111-1; Viral Vector Facility, Zurich University) were applied at DIV1 on top of the hippocampal CA1 region by a microinjector (Eppendorf, FemtoJet) aided by a stereoscopic microscope (Leica, M80). This resulted in widespread, but sparse GFP and Gs-HA expression in GABAergic neurons, which partially overlapped. Two-photon time lapse imaging was performed when slices were kept 2 to 3 weeks *in vitro*. After a baseline period (5 time points), Gs signaling was activated by bath application of 10 µM CNO (Tocris Bioscience), which was continued until the end of the experiment. P*ost hoc* immunostaining was performed using rat anti-HA primary antibodies (Roche, #11 867 423 001) and anti-rat Alexa568 (Life Technologies, A11077, 1:500) as secondary antibodies. We selected slices with good GFP labeling in the dendritic layer for the two-photon experiments and in 4 out of 13 slices we were able to identify up to 5 axons of each type within the imaging area. Identification of HA+ and HA-axons was performed in Fiji, bouton dynamics analysis in Matlab as described above.

### Statistical Analysis

All experiments were performed and analyzed blindly. Live imaging experiments for bouton dynamics analysis were performed in paired slices from the same animal and the same culture. Statistical analysis was performed using GraphPad Prism software. Data are reported as mean ± standard error unless stated otherwise. The variance between axons was larger than the variance between slices, indicating that individual axons are independent measurements. Results from treatment and control experiments were compared using the nonparametric Mann-Whitney U test (MW). Distributions were compared with Kolmogorov-Smirnov test (KS). Multiple comparisons were made using two-way ANOVA (2w ANOVA) followed by Sidak’s test. Repeated two-way ANOVA analysis was used for comparing NP bouton density and NP presence over time. P-values (not adjusted for multiplicity) are indicated in the figure legends. Differences were considered significant when *p* < 0.05 (\**p* < 0.05, \*\**p* < 0.01, \*\*\**p* < 0.001).

## Acknowledgements

This work was supported by a CSC scholarship (JL) and by the Netherlands Organisation for Scientific Research, as part of the research program of the Foundation for Fundamental Research on Matter (FOM) (DLHK; #15PR3178). We thank Lotte Herstel for helping with experiments in Fig. 1E-F and René van Dorland for excellent technical support.

## Author contributions

JL, DLHK, CJW designed the experiments. JL, DLHK, MR, BJBV, ACJV, FB performed the experiments. JL, BJBV, ACJV, FB, MSG, MR analyzed the data. CJW wrote the manuscript with input from all other authors.

## Conflict of interest

The authors declare no competing financial interests.

## Notes

### Competing Interest Statement

The authors have declared no competing interest.

